# Functional neural architecture of cognitive control mediates the relationship between individual differences in bilingual experience and behaviour

**DOI:** 10.1101/2022.10.06.510955

**Authors:** Felix Carter, Vincent DeLuca, Katrien Segaert, Ali Mazaheri, Andrea Krott

**Author notes:** Corresponding author: Felix Carter, School of Psychology, University of Bristol, Bristol BS8 1TU, United Kingdom. **Author contributions**: AK, KS, and AM conceived the study, FC and VD collected the data, all authors contributed to the design of the analysis, FC and VD performed the analysis, FC and AK wrote the manuscript with contributions of AM, KS, and VD. **Classification:** Social Sciences (Psychological and Cognitive Sciences), Biological Sciences (Neuroscience).

## Abstract

Bilinguals have often, but not always, been found to outperform monolinguals on domain-general attentional control. Inconsistent findings have been argued to stem, at least partly, from treating bilingualism as a uniform category and from not considering how neural adaptations to bilingual experiences modulate behavioural outcomes. The present study investigated how patterns of language experience, including language switching behaviour, duration and intensity/diversity of bilingual language use, influence the brain processes underlying cognitive control, and how these in turn translate to cognitive control performance. We examined reaction times and spectral dynamics of the electroencephalograms (EEG) of two-hundred-and-thirty-nine participants (about 70% bilinguals) with diverse language experiences during two cognitive control paradigms testing interference suppression (flanker and Simon task). Using structural equation modelling, we found that different bilingual experience factors were related with neurocognitive measures, which in turn were related with behavioural interference effects, for the flanker but not the Simon task. More specifically, increased frequency of language switching and intensity / diversity of bilingual language usage was negatively related to induced top-down control measures (especially midline-frontal theta), which in turn was beneficial for interference control. In contrast, duration of bilingual engagement correlated negatively with evoked bottom-up control measures (especially P3) and was therefore detrimental to interference control. We demonstrate here for the first time how the different factors of bilingual experience lead to different neural adaptations which impact behavioural outcomes.

**Significance statement:** Like other intensive experiences, bilingualism leads to brain adaptations. It results in structural changes in language areas, and, due to demands on language control, in brain areas associated with domain-general cognitive control. Related to this, bilinguals often outperform monolinguals on cognitive control tasks. But what is often ignored is that bilingualism is a multi-dimensional phenomenon, with variations such as diversity of language usage and duration of language use. The present large-scale study of neural functioning in bilingualism revealed for the first time how individual differences in bilingual experience lead to adaptations to brain functioning which in turn affect cognitive control behaviour. It exemplifies how the complexity of individual experiences plays a fundamental role in brain function.

## Introduction

Bilingualism has been observed to confer adaptations in domain-general attentional control (Bialystok, 2017; Bialystok & Craik, 2022; Bialystok, Craik, & Luk, 2012). Managing more than one language, as bilinguals do to varying degrees in everyday life, requires not only language specific but also a multitude of domain-general cognitive functions (Anderson, Chung-Fat-Yim, Bellana, Luk, & Bialystok, 2018; De Baene, Duyck, Brass, & Carreiras, 2015; Garbin et al., 2010). Bilingual speakers need to inhibit the non-target language to avoid language intrusion and mixing, given that their languages are constantly and simultaneously active (Kaushanskaya & Marian, 2007; Kroll, Dussias, Bice, & Perrotti, 2015). This continuous demand on cognitive control to manage one’s languages has been argued to train domain-general control mechanisms. Evidence for this assertion has been found with various executive function tasks, in which bilinguals often outperform a matched group of monolingual speakers (see overviews in Bialystok, 2017; Bialystok & Craik, 2022; Bialystok et al., 2012). This enhanced performance has historically been dubbed the ‘bilingual advantage’.

However, the observation of the ‘bilingual advantage’ has not been unequivocal, with some studies reporting inconsistent and sometimes seemingly contradictory findings (for recent meta-analyses see Donnelly, Brooks, & Homer, 2019; John G. Grundy, 2020; Lehtonen et al., 2018; van den Noort et al., 2019). These inconsistencies have been suggested to emerge, at least partly, from treating bilingualism as a categorical variable (e.g., Luk & Bialystok, 2013; Surrain & Luk, 2019; H. Yang, Hartanto, & Yang, 2016) and from not considering the potentially modulatory role of any neural adaptations underlying these cognitive outcomes (see e.g., Abutalebi et al., 2012; Ansaldo, Ghazi-Saidi, & Adrover-Roig, 2015).

There has recently been an increase in investigations attempting to link variations in behavioural performance in tasks employing attentional control, and patterns of neurocognitive outcomes respectively, with differences in bilingual experiences such as intensity of bilingual engagement or age of acquisition of a second language (see for review DeLuca, Segaert, Mazaheri, & Krott, 2020). While these studies are very informative, few to no studies to date have directly examined the link between neural and behavioural outcomes and how these relationships are modulated by patterns of language experience. Specifically, little is known about how patterns of language experience (such as the above) influence the brain processes underlying cognitive control and crucially how these in turn translate to cognitive control performance. Of particular interest is the proposal that increasing bilingual language usage leads to a shift from anterior cortical control mechanisms to subcortical control mechanisms which is proposed as an adaptation towards increased efficiency in handling control demands (see e.g., Grundy et al., 2017; Pliatsikas, 2020).

Electrophysiological (EEG) studies focusing on evoked response markers of bilingual effects in interference suppression, as in the flanker and Simon tasks, have investigated modulations of the N2 and P3 components (see also investigations into the N450 in Stroop tasks by Coderre & van Heuven, 2014; Heidlmayr, Hemforth, Moutier, & Isel, 2015). Importantly for our study, Kousaie and Phillips (2012) recorded EEG during Stroop, Simon, and flanker task performance in monolingual and bilingual young adults. While there were no behavioural differences between the two groups for any of the tasks, there were ERP differences, albeit not consistent across tasks. In the Stroop task, bilinguals had reduced N2 amplitudes and P3 latencies across all experimental conditions. For the Simon task, bilinguals had smaller P3 amplitudes. The flanker task was the only task that showed group differences in the interference effect, with monolinguals showing a larger interference effect in P3 peak latencies. Kousaie and Phillips (2017) conducted the same study with older adults and found similar results.

In addition to the evoked responses, oscillatory dynamics provide further insight into the neurocognitive underpinnings of interference control, with midfrontal theta (3-7 Hz), alpha (8-12 Hz), as well as beta (15-30 Hz) oscillations playing distinct roles. Midfrontal theta plays a critical role in cognitive control (e.g., Luu et al., 2004; Trujillo & Allen, 2007; Cavanagh et al., 2009). Nigbur, Ivanova and Stuermer (2011) found increased theta power over midfrontal sites when responses were successfully inhibited in incongruent trials (relative to congruent trials) for both a flanker and Simon task. Theta power reflects recruitment of additional cognitive control (Cavanagh & Frank, 2014; Sauseng, Hoppe, Klimesch, Gerloff & Hummel, 2007) and as such is examined in the current study as a neurocognitive marker of proactive control mechanisms for interference suppression.

Alpha oscillations have functionally been related to cortical inhibition (Hanslmayr et al., 2007; Lange et al, 2013; Romei et al, 2008; van Dijk et al, 2008; Zumer et al, 2014) and have been shown to enhance feature-based attention (van Diepen, Miller, Mazaheri & Geng, 2016). As such, alpha dynamics are likely a crucial component of interference suppression. The role of beta activity in interference control is more ambiguous. Beta power over the motor cortex is suppressed just prior and during a voluntary movement (Jasper & Penfield, 1949; Salmelin & Hari, 1994a; Pfurtscheller et al., 2003; Jurkiewicz et al., 2006). Within the context of interference control, previous work has suggested that a beta increase following a motor response (i.e., beta rebound) reflects the brain processes involved in maintaining the ‘status quo’ (“maintenance of the sensorimotor set”) after interference suppression (Engle & Fries, 2010). To our knowledge, few to no studies to date that have examined potential variations in oscillatory dynamics underlying differences in interference suppression between monolinguals and bilinguals (but see Calvo & Bialystok, 2021 for an example of oscillatory dynamics pertaining to bilingual effects on proactive interference measures of attentional control).

In the present study, we aimed to determine how variations in bilingual experience impact the neural architecture underlying attentional control and how the latter corresponds to variations in behavioural performance. We examined spectral dynamics of the EEG (both evoked and induced) of a large sample of participants with diverse language experiences during two distinct cognitive control paradigms that are typically used in this area of research and that have revealed superior performance of bilingual speakers compared to monolingual speakers, the flanker and Simon task (see overview in Zhou & Krott, 2016). Beyond a traditional analysis of group level differences between monolingual and bilingual participants, we aimed to investigate effects of specific language experiences, that is to examine bilingualism as a continuum of experiences. We used the Unified Bilingual Experience Trajectory (UBET) framework (DeLuca et al., 2020) as a basis for our predictions and experience measures. The UBET framework is based on previous findings and models, specifically the Adaptive Control Hypothesis (ACH) (Abutalebi & Green, 2016; Green & Abutalebi, 2013), the Conditional Routing Model (CRM) (Stocco et al., 2014; Stocco, Lebiere, & Anderson, 2010), the Bilingual Anterior to Posterior and Subcortical Shift (BAPSS) framework (Grundy et al., 2017) and the Dynamic Restructuring Model (DRM) (Pliatsikas, 2019b). These models stress different aspects of bilingualism and how these affect various brain areas involved in cognitive control. Combining and extending the predictions from these models, the UBET framework distinguishes four general aspects of bilingual experience: diversity/intensity of bilingual engagement (see also the ACH), relative language proficiency (see CRM), duration of bilingual engagement (see also BAPSS, DRM), and language switching (see CRM). The UBET framework attempts to account for effects of these language experiences on both structural and functional brain adaptations. We focus herein on functional adaptations captured by electrophysiological markers of control. Specifically, we investigated the relationship between the different dimensions of the bilingual experience of the UBET framework on both top-down and bottom-up cognitive control, measures by oscillatory and evoked brain responses respectively, and crucially how the two types of control might variably impact behavioural performance in the two cognitive control paradigms. Given that both tasks have been argued to measure inhibitory control and given that they have shown very similar effects within the same population (Zhou & Krott, 2018), we expected similar neural processes to govern behavioural effects in both tasks.

## Results

### Monolinguals respond faster, but no group speed difference in interference effect

We analysed reaction times (RTs) for both the flanker and Simon task with linear mixed effect models using the *lmer* function of the lme4 package (Bates et al., 2015) in R version 4.1.2 (R Core Team, 2021). The models contained the fixed factors language group (monolinguals versus bilinguals), condition (congruent versus incongruent) and their interaction, as well as random intercept plus random slopes for participants^1^.

In line with previous literature (see Lu & Proctor, 1995; Ridderinkhof, Wylie, van den Wildenburg, Bashore & van der Molen, 2021 for reviews of the Simon and flanker effects respectively), we observed a significant congruency effect on RTs in both the flanker (n = 227, 162 bilinguals, t(224.7) = 48.83, partial η^2^ *=* 0.91 *p* < 0.001) and Simon task (n = 207, 148 bilinguals, t(204.6) = 15.98, partial η^2^*=* 0.56, *p* < 0.001). As expected, for the flanker task, RTs were faster for congruent (M = 474.64ms, SD = 98.8) than incongruent (M = 587.49ms, SD = 109.28) trials. Similarly, in the Simon task, RTs were faster for congruent (M = 436.37ms, SD = 121.2ms) than incongruent (M = 460.57 SD = 110.2) trials.

We also found a global speed advantage for monolinguals for both the flanker (t(226.8) = 3.47, partial η^2^ *=* 0.05, *p* < 0.001) and Simon task (t(206.78) = 2.23, partial η^2^*=* 0.02, *p* = 0.03), without group differences in interference effects. Thus, monolinguals were significantly faster to respond irrespective of trial type. In the flanker task, the mean RT for monolinguals was 481.73ms (SD = 107.52) compared to 509.65ms (SD = 113.15) for bilinguals. However, the language group x condition interaction was not significant t(224.7) = 0.65, *p* = 0.52). Likewise, in the Simon task, the overall RTs for monolinguals was 436.42ms (SD = 114.1) compared to 453.13ms for bilinguals (SD = 117.1ms). Again, the language group x condition interaction was not significant, t(204.6) = 0.27, *p* = 0.79.

In sum, behavioral results show previously reported interference effects in both tasks. In addition, bilinguals responded more slowly than monolinguals in the two tasks, without any group differences in interference effects.

### Bilinguals show attenuated and later evoked EEG responses as well as smaller interference effect on P3 amplitudes

Stimulus-locked ERP waveforms were analysed using mixed analyses-of-variance (MANOVAs) for mean amplitude and peak latency of each component (N2 and P3). MANOVAs included condition (congruent versus incongruent) as a within-subject factor, language group (bilingual versus monolingual) as a between-subject factor and the condition x language group interaction. Latency ranges and channel locations were located based on previous studies (see Patel & Azzam, 2005, for a review) and confirmed with visual inspection of the grand-averaged waveforms. N2 was measured at FCz, from 375-425ms (Flanker task) & 275-325ms (Simon task). P3 was measured at Pz from 300-650ms (both tasks).

Stimulus-locked ERP waveforms are shown in Figure 1 and N2 and P3 amplitudes and latencies are listed in Table 1. For both tasks, and in line with what had previously been reported in the literature (e.g., Bartholow et al, 2005; Heil, Osman, Wiegelmann, Rolke & Hennighausen, 2000; Kopp, Rist & Mattler, 1996; Yeung, Botvinick & Cohen, 2004; Xie, Cao, Li & Li, 2020), we observed congruency effects (i.e., amplitudes being larger for incongruent than congruent trials) for P3 in both tasks and for N2 in the Simon task

We also found that, compared to monolinguals, bilinguals had attenuated N2s. Exclusive to the flanker task, they also had attenuated P3s and a smaller P3 congruency effect. For the flanker task, N2 amplitudes did not differ between conditions, *F*(1, 237) = 0.003, partial η^2^< 0.001*, p* = 0.960, but was significantly more pronounced for monolinguals compared to bilinguals, *F*(1, 237) = 4.93, partial η^2^*=* 0.02, *p* = 0.027. Also, N2 peak latencies occurred significantly later for bilinguals compared to monolinguals, *F*(1, 237) = 5.44, partial η^2^*=* 0.02, *p* = 0.021. P3 amplitudes were significantly larger for incongruent than congruent trials, *F*(1, 237) = 33.25, partial η^2^*=* 0.12, *p* < 0.001, and for monolinguals compared to bilinguals, *F*(1, 237) = 7.27, partial η^2^ *=* 0.03, *p* = 0.008. Here, the language group x condition interaction was significant, *F*(1, 237) = 17.97, partial η^2^*=* 0.07, *p* < 0.001, with the interference effect being smaller in bilinguals than monolinguals. Furthermore, P3 amplitudes peaked significantly later for incongruent than congruent trials, *F*(1, 237) = 255.83, partial η^2^ *=* 0.52, *p* < 0.001, and for bilinguals compared to monolinguals; *F*(1, 237) = 5.49, partial η^2^ *=* 0.02, *p* = 0.02.For the Simon task, N2 amplitudes were significantly more pronounced for incongruent compared to congruent trials, *F*(1, 206) = 22.31, partial η^2^ *=* 0.1 *p* < 0.001, and for monolinguals compared to bilinguals, *F*(1, 206) = 14.75, partial η^2^*=* 0.07, *p*< 0.001. N2 amplitudes also peaked significantly later for incongruent compared to congruent trials, *F*(1, 206) = 4.996, partial η^2^ *=* 0.02, *p* = 0.027. P3 amplitudes were significantly larger for incongruent than congruent trials (*F*(1, 206) = 25.92, partial η^2^*=* 0.12, *p* < 0.001), and peaked significantly later for incongruent compared to congruent trials, *F*(1, 207) = 57.19, partial η^2^ *=* 0.22, *p* < 0.001^2^.

In sum, results for both tasks replicated previously reported congruency effects on P3 (and N2 for the Simon task). Compared to monolinguals, bilinguals exhibited attenuated and later N2 and P3 responses in the flanker task as well as attenuated N2 responses in the Simon task. Importantly, bilinguals also showed a smaller interference effect on P3 amplitudes in the flanker task.

**Table 1:** Mean amplitudes and peak latencies for N2 & P3 components for both flanker and Simon task.

| **Flanker task** | N2 Amplitude (SD) | N2 Latency (SD) | P3 Amplitude (SD) | P3 Latency (SD) |
| --- | --- | --- | --- | --- |
| *Congruent trials* |  |  |  |  |
| Monolingual | -1.91mv (2.31) | 384.54ms (31.17) | 3.58mv (1.56) | 393.18ms (66.03) |
| Bilingual | -1.16mv (1.84) | 391.76 (31.16) | 3.13mv (1.81) | 423.93ms (72.86) |
| Groups Collapsed | -1.38mv (2.01) | 389.62ms (31.27) | 3.26mv (1.75) | 414.80ms (72.15) |
| *Incongruent trials* |  |  |  |  |
| Monolingual | -1.92mv (2.35) | 386.62ms (30.09) | 4.62mv (2.07) | 494.51ms (87.49) |
| Bilingual | -1.37mv (1.87) | 394.21ms (27.32) | 3.60mv (2.31) | 504.70ms (87.32) |
| Groups Collapsed | -1.53mv (2.03) | 392.03ms (28.32) | 3.91mv (2.29) | 501.67ms (87.31) |
| **Simon task** |  |  |  |  |
| *Congruent trials* |  |  |  |  |
| Monolingual | -3.00mv (2.06) | 302.47ms (18.31) | 3.84mv (1.48) | 363.23ms (54.71) |
| Bilingual | -1.67mv (2.04) | 302.03ms (19.31) | 3.35mv (1.98) | 375.92ms (67.32) |
| Groups Collapsed | -2.05mv (2.13) | 302.15ms (18.99) | 3.49mv (1.86) | 372.26ms (64.07) |
| *Incongruent trials* |  |  |  |  |
| Monolingual | -3.20mv (2.07) | 304.20ms (17.29) | 4.04mv (1.44) | 394.20ms (72.78) |
| Bilingual | -2.01mv (1.97) | 305.01ms (18.62) | 3.48mv (2.08) | 410.16ms (81.56) |
| Groups Collapsed | -2.36mv (2.07) | 304.78ms (18.21) | 3.64mv (1.93) | 405.58ms (79.28) |

**Figure 1:**
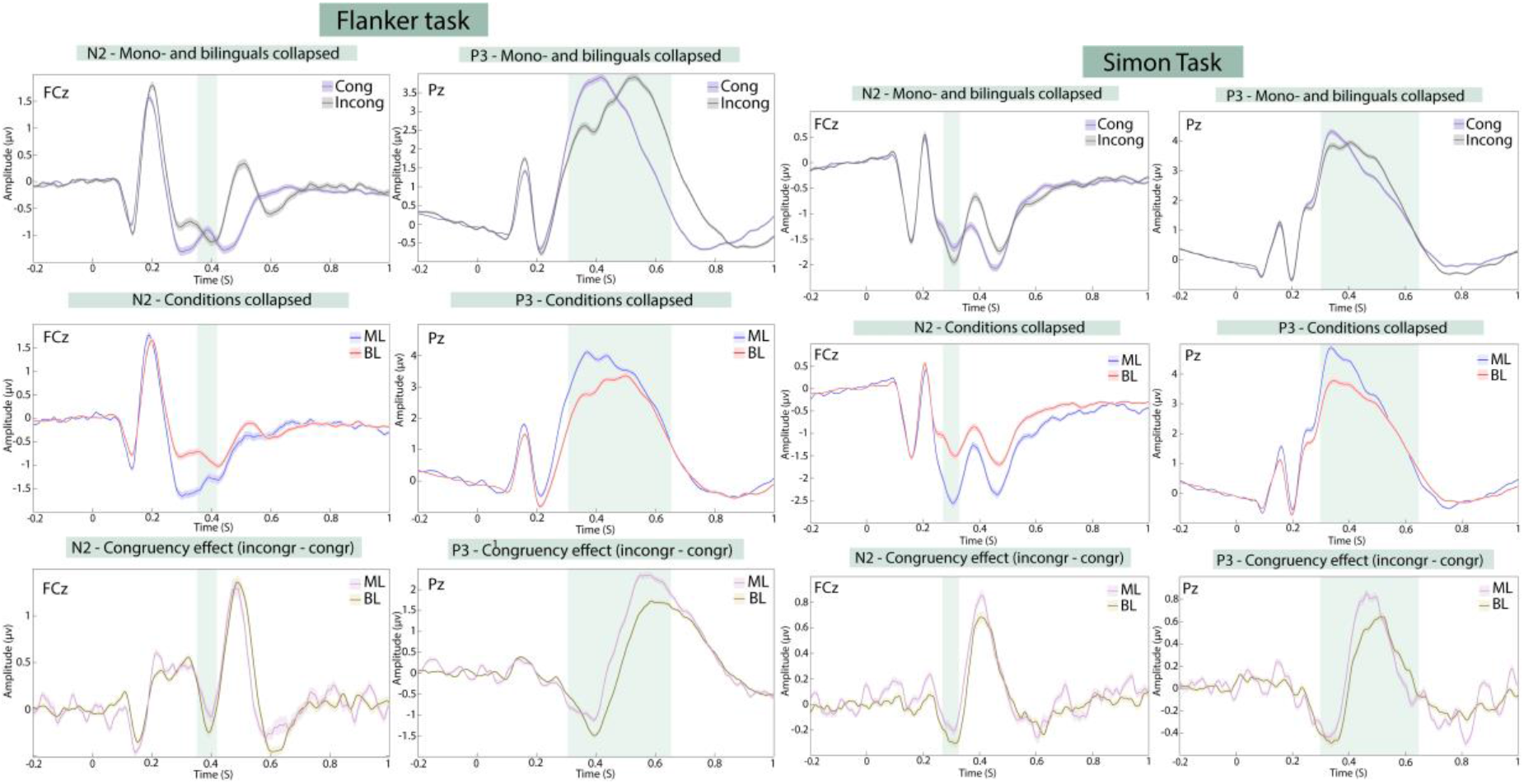
Bilinguals exhibited attenuated and delayed N2 and P3 components. Stimulus-locked ERP waveforms with grand-averaged N2 (left columns) and P3 (right columns) for flanker task (upper panel) and Simon task (lower panel). For each task, shown are averaged data by condition (upper row), by group (middle row) and congruency effects (i.e., condition differences; bottom row). Shaded coloured regions represent standard errors; shaded boxes indicate latency ranges analysed.

### Congruency effect differences between language groups were not consistent between the two cognitive control tasks

All analyses reported in this section are conducted on induced spectral activity, which was obtained by subtracting the evoked activity (i.e., the time-frequency response of the averaged time-locked activity) from the time-frequency analysis output for each participant. Non-parametric cluster-based permutation testing (Maris & Oostenveld, 2007) was used to correct for multiple comparisons, with clusters defined as two or more channel pairs whose t-test significance was *p* < 0.05. The test statistic was defined as the maximum of cluster-level statistics obtained from summing t-values within each cluster. The reference distribution was approximated by 1000 Monte Carlo simulations.

The components of interference control were identified by collapsing both language groups and using a cluster-based permutation test to find the points of maximal difference between conditions. For each task, induced theta (4 – 7 Hz) and alpha (8 – 12 Hz) activity were compared between conditions between 0 and 1000ms post-stimulus using dependent samples t-tests. Figures 2 and 3 show the TFRs of power locked to the onset of the stimuli for the flanker and Simon tasks, respectively. In line with previous literature (e.g., Cavanagh & Frank, 2014; Duprez, Gulbinaite & Cohen, 2020), we found that incongruent stimuli across both tasks induced significantly greater theta power (Figures 2A and 3A) than congruent stimuli (p<0.05, Monte Carlo P value). As shown in Figures 2C and 3C, this congruency effect had a maximal distribution over midline electrodes. The maximal difference in theta power appeared 250-750ms post-stimulus for the flanker task and 100-500ms post-stimulus for the Simon task.

In terms of group differences, we found divergent patterns for the two tasks. For the flanker task, we found that the monolinguals had a significantly larger theta increase induced by incongruent trials than the bilinguals (p<0.05, Monte Carlo P value; see Figure 2B). This difference in theta power appeared more anterior than the congruency effect observed between conditions (Figure 2F). Interestingly, in the Simon task it was the bilinguals which had the larger theta increase (Figure 3B) for incongruent stimuli (p<0.05, Monte Carlo P value). The difference in theta power between the language groups, shown in Figure 3D, appeared spatially more widespread than the congruency effect in theta observed between conditions (Figure 3C).

Furthermore, we found differences in alpha suppression, but only in the flanker task. Incongruent flanker stimuli induced significantly greater alpha suppression than congruent flanker stimuli (p<0.05, Monte Carlo P value), which appeared maximal over posterior sites from 600-1000ms post-stimulus (see Figure 2D). This effect was not found to be different for the two language groups or present in the Simon task.

Finally, we observed differences in beta rebound, but again only for the flanker task. Incongruent flanker stimuli induced significantly greater beta rebound than congruent flanker stimuli (p<0.05, Monte Carlo P value), with the maximal difference over frontocentral sites from 1100-1500ms post-stimulus (see Figure 2E). There was also a group difference with regards to the interference effect. Monolinguals displayed a significantly larger beta increase induced by incongruent flankers than the bilinguals (p<0.05, Monte Carlo P value), with this difference being most apparent at frontal and occipitoparietal sites.

In sum, we found previously reported congruency effects for theta and alpha plus a congruency effect for beta rebound, with the effects in alpha and beta only for the flanker task. There were also group differences in congruency effects in theta and beta, with the effect in theta having opposite patterns across the two tasks. While monolinguals showed a larger effect in the flanker task, they showed a smaller effect in the Simon task. In addition, these group differences had different distributions compared to the condition differences. The effect in the flanker task was more anterior and the one in the Simon task was more widespread. In terms of beta, monolinguals displayed a significantly larger congruency effect in beta rebound than bilinguals in the flanker task.

**Figure 2:**
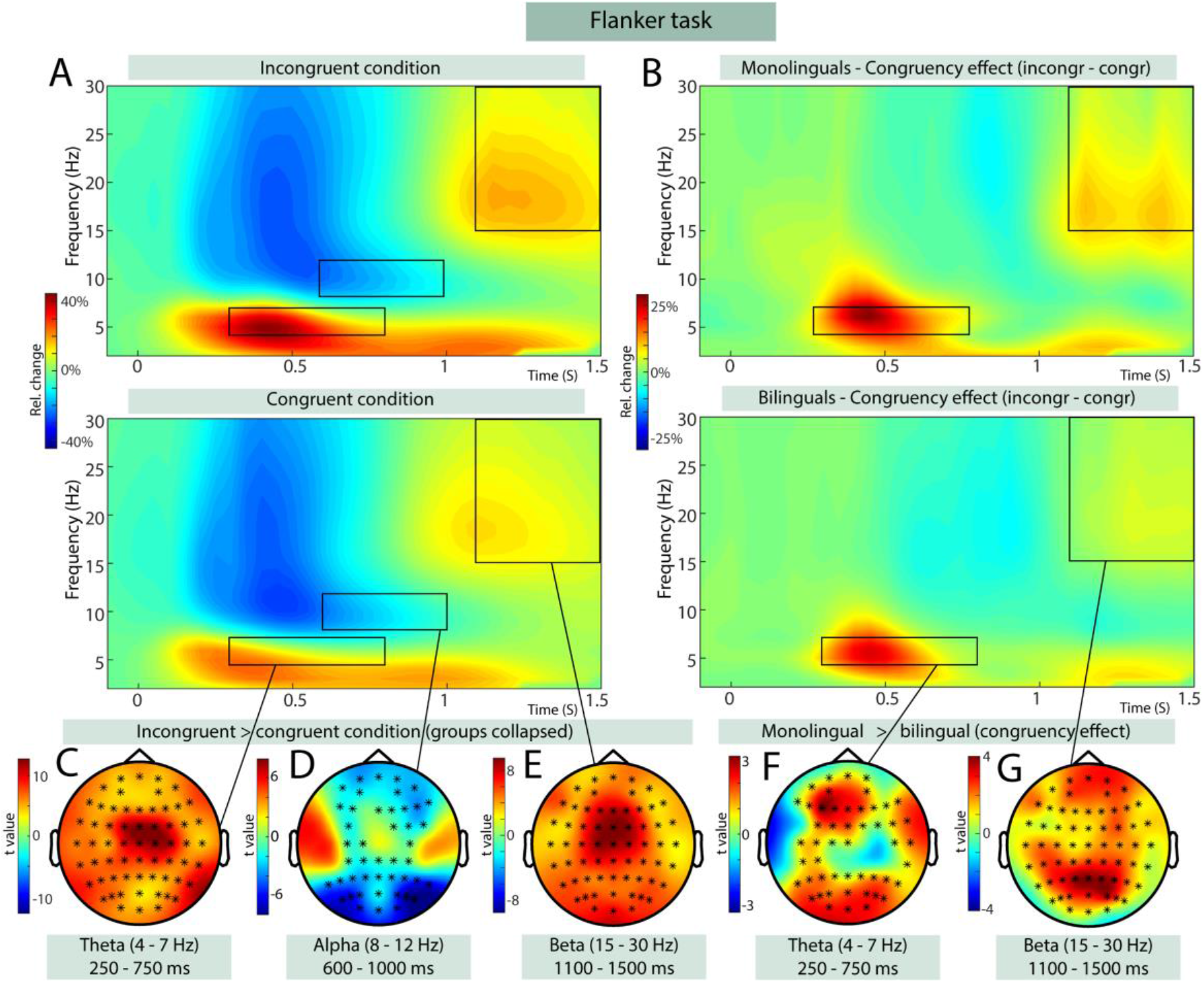
In the Flanker task, monolinguals had a significantly larger theta increase induced by incongruent trials than bilinguals. Figure shows TFR of power (relative to pre-stimulus baseline) locked to stimulus onset during the flanker task. A) TFRs collapsed across language groups for congruent and incongruent trials, B) congruency effects (incongruent-congruent trials) for bi- and monolinguals, C-E) scalp topographies illustrating the clusters of electrodes that show the most pronounced mean congruency effect for theta (C), alpha (D) and beta (E) activity respectively, F-G) scalp topography illustrating the clusters of electrodes that show the most pronounced difference in congruency theta (F) and beta (G) activity between bilinguals and monolinguals.

**Figure 3:**
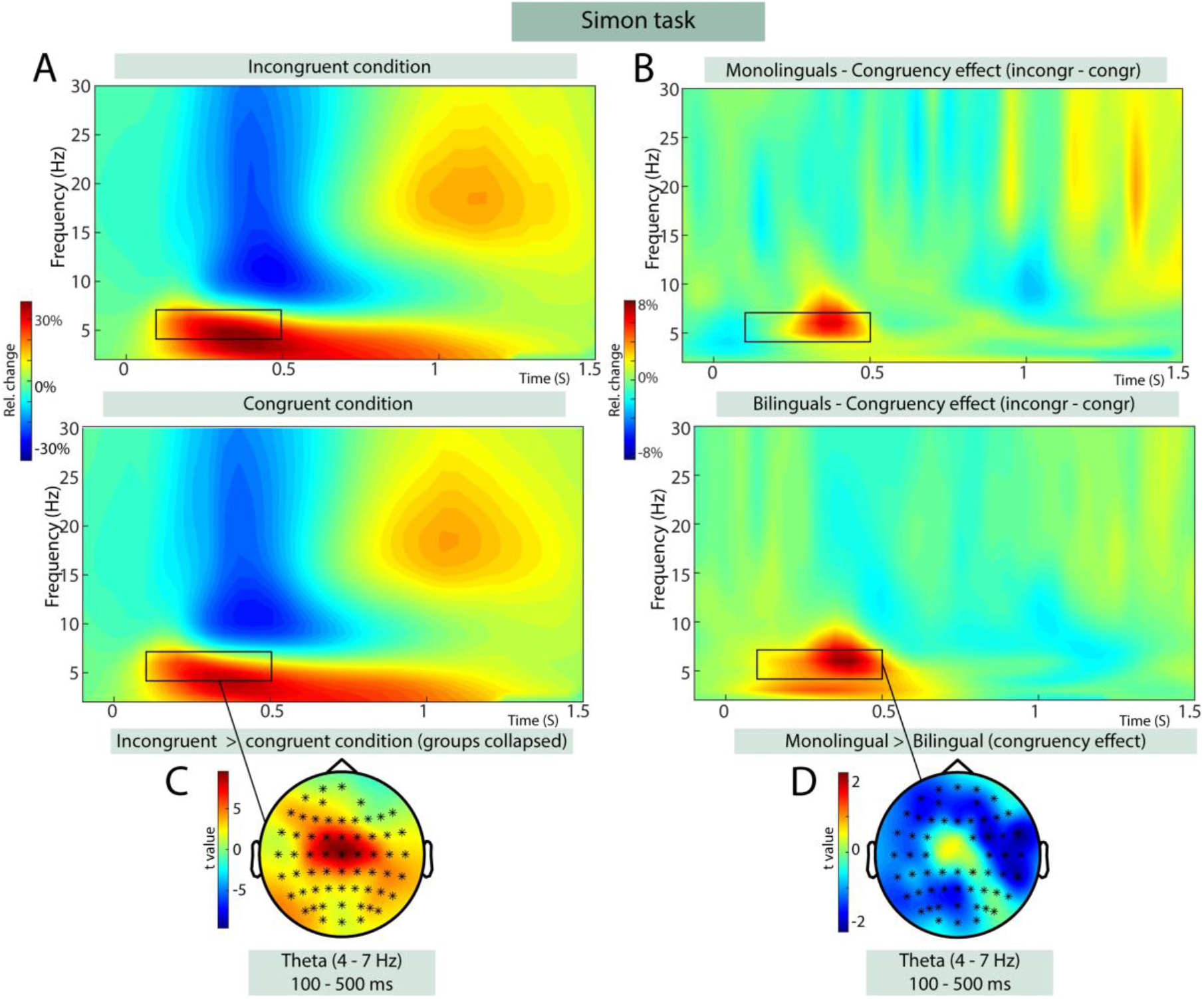
During the Simon task, it was the bilinguals which had the larger theta increase in the incongruent trials. Figure shows TFR of power (relative to pre-stimulus baseline) locked to stimulus onset during the Simon task. A) TFRs collapsed across language groups for congruent and incongruent trials, B) congruency effects (incongruent-congruent trials) for bi- and monolinguals, C) scalp topography illustrating the clusters of electrodes that show the most pronounced mean congruency effect for theta activity, D) scalp topography illustrating the clusters of electrodes that show the most pronounced difference in congruency theta between bilinguals and monolinguals.

#### Operationalising bilingual experience

In order to assess the bilingual experience of our participants we utilised a number of established questionnaires (see methods) which focused on: the extent of non-English language use and its duration as well as the degree of language switching, and proficiency. Using confirmatory factor analysis (CFA), the responses of questionnaires and proficiency measures were loaded onto 4 latent variables to test the effect of bilingual experience on interference suppression (derived from the UBET model; DeLuca et al, 2020). Trelhese were *duration of bilingual language use*, *intensity and diversity of language use* (hereafter *intensity/diversity)*, *language switching* and *relative proficiency.* The CFA was performed using the *lavaan* package in R, with variables loaded as described in the methods section and presented in Table 2. A value of <0.08 was used as an acceptable value for Root Mean Square Error of Approximation (RMSEA) and Standardized Root Mean Square Residual (SRMR; Browne & Cudeck, 1993). A value of >0.9 was used as an acceptable value for the Comparative Fit Index (CFI; Bentler, 1990) and Tucker-Lewis Index (TLI; Tucker & Lewis, 1973).

Model fit indices for the CFA indicated acceptable goodness-of-fit for the four-factor model with the factors of i) duration ii) intensity/diversity iii) language switching, and iv) relative proficiency (Comparative Fit Index (CFI) = 0.916; Tucker-Lewis Index (TLI) = 0.906; Root Mean Square Error of Approximation (RMSEA) = 0.074; Standardized Root Mean Square Residual (SRMR) = 0.075). Verbal fluency measures were dropped from the final CFA due to a large number of missing values (>10% of total data).

Factor scores were calculated from the CFA output using the lavPredict function in the *lavaan* package (Rosseel et al., 2012). These factor scores were subsequently entered into a structural equation model (SEM) with the behavioral performance and EEG measures. The *relative proficiency* latent variable was removed from the SEM due to model non-convergence, caused by high correlation of error variance with the *intensity/diversity* variable.

### Bilingual experience predicts neural correlates of interference suppression which in turn predict behavioural response patterns

Based on our proposed UBET model we hypothesised that increased *diversity/ intensity of bilingual language use* and increased *language switching* would create enhanced *EF demands,* which would in turn reduce reliance on cortical *top-down control* mechanisms for managing interference, reflected in reduced congruency effects in induced frontal theta, posterior alpha power, and midfrontal beta power. *Duration of bilingual language use* was hypothesised to affect the *efficiency* of interference resolution, which in turn would affect *bottom-up control*, reflected in how quickly conflict is detected (represented by N2 latency) and resolved (represented by P3 latency) and the resources allocated to detecting and resolving interference (represented by N2 and P3 amplitude, respectively). EEG measures were hypothesised to jointly predict the behavioural congruency effect; thus, congruency effects in evoked and induced neural activity were both regressed onto the RT congruency effect.

Modelling the two tasks in a single model led to poor model fits. We therefore modelled them separately. Figure 4 shows the results of the structural equation modelling for both tasks. For the flanker task, the model resulted in acceptable fit (CFI = 0.929; TLI = 0.902; RMSEA = 0.058; SRMR = 0.076). P3 amplitude (*p <* 0.001) and latency (*p<* 0.001) loaded significantly onto the *evoked neural activity* factor, while N2 latency and amplitude did not (*p*s > 0.05). Only the congruency difference in induced frontal theta power significantly loaded onto the *induced neural activity* factor (*p =* 0.001), although the loading of beta power was marginal (*p =* 0.07). All regression paths of the model were significant (all *p*s < 0.05).

In contrast, for the Simon task data, only theta and beta power significantly loaded onto latent factors, and no regression paths were significant (all *p*s > 0.05). Possible explanations for the apparent discrepancy of the models for the two tasks are considered in the *Discussion* section.

These results support the hypothesis that bilingual experience leads to functional neurocognitive adaptations which are observable during an interference suppression (flanker) task, and that these adaptations themselves impact behavioural indicators of interference suppression.

**Figure 4:**
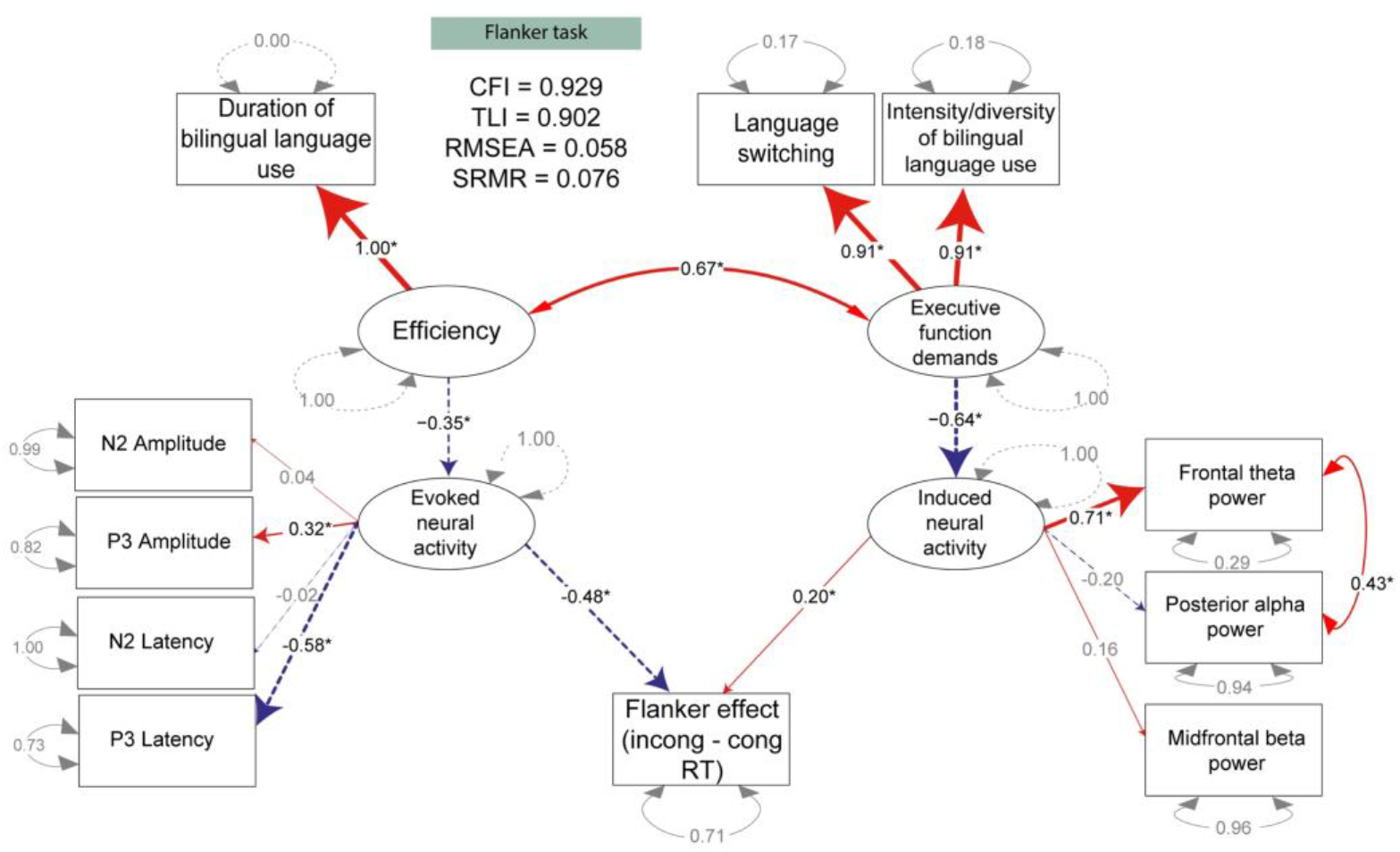
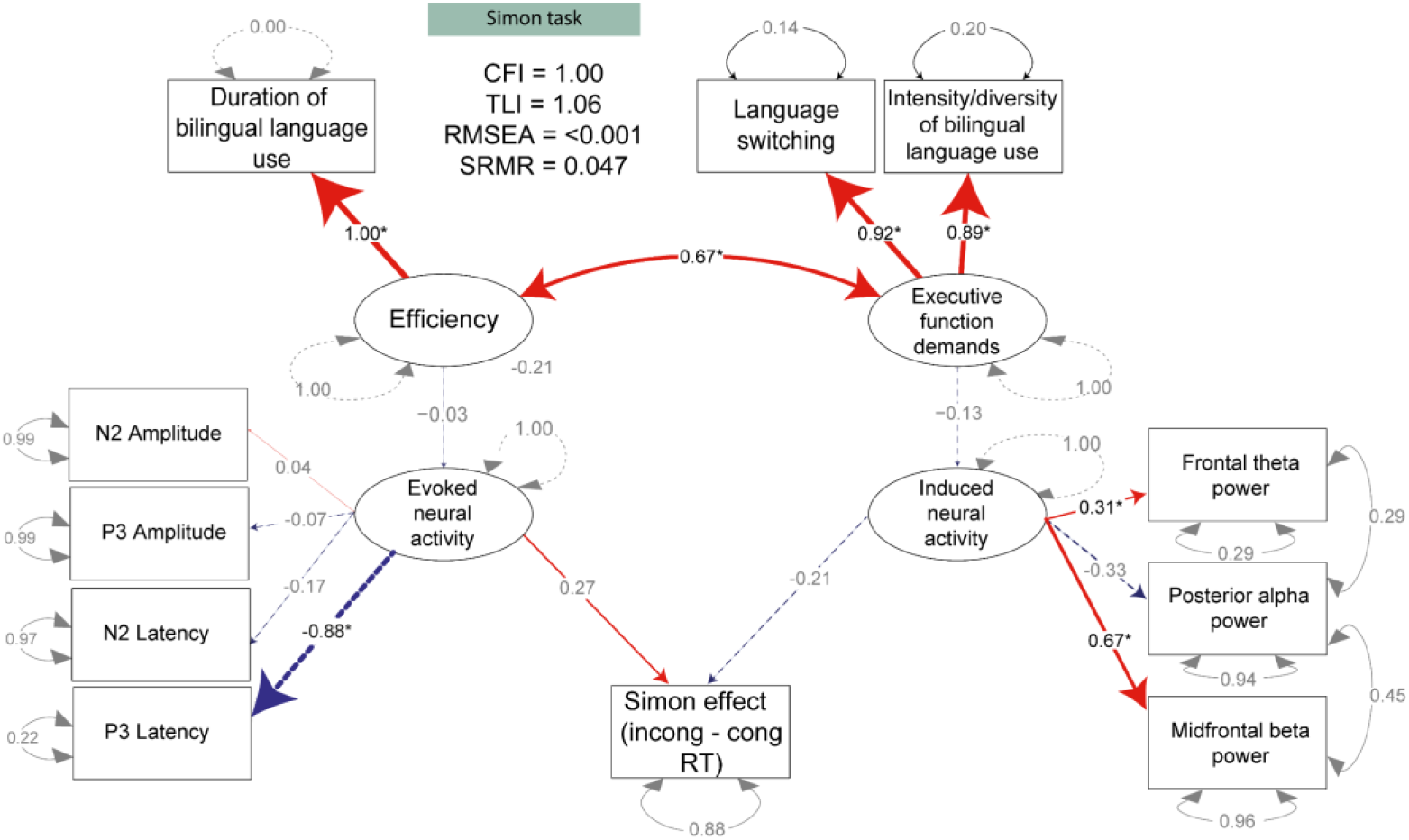
SEM plots for Flanker (above) and Simon (below) tasks. Latent variables are indicated by ovals. Observed variables are indicated by rectangles. Straight arrows indicate regression relationships or factor loadings. Curved arrows indicate correlated error variances. Significant (p < 0.05) factor loadings or regression relationships are marked by *. Numbers next to rounded arrows are covariance coefficients for residual errors (dotted rounded lines show single indicator variables). Numbers next to straight lines are standardized path coefficients (regression weights) or factor loadings.

## Discussion

In the current investigation we examined how the different elements of the bilingual experience impacted the neural architecture underlying attentional control and how the latter corresponds to variations in behavioural performance in two interference suppression tasks. We found that both increased frequency of language switching and intensity / diversity of bilingual language usage corresponded to diminished interference effects in induced top-down control measures (theta power) in the flanker task which in turn was beneficial for interference control. In other words, frequent engagement with both languages and frequent switching between languages seemed to have trained executive control, which meant that proactive top-down control was more efficient and thus led to less interference of flankers. In contrast, duration of bilingual engagement correlated negatively with interference effects in evoked bottom-up control measures (specifically P3) which was in turn was detrimental to interference control. Thus, having used two languages for longer meant that attentional resources for interference suppression were reduced and delayed, which meant that interfering flankers led to increased response delays.

Our findings broadly support the hypotheses of the UBET framework (DeLuca et al., 2020) and the models it is based on, which detail the structural and functional neurocognitive adaptations to the cognitive control demands associated with bilingual engagement.

Crucially, as we discuss below, these neural adaptations to experience have implications for attentional control processes.

Previous work has suggested that, with time, bilingual engagement leads to a shift from a reliance on anterior cortical control regions to posterior subcortical regions (see also the BAPSS framework: Grundy et al., 2017; and the DRM: Pliatsikas, 2020). Our EEG results for the flanker task are in line with this hypothesis. Bilinguals exhibited reduced neural responses associated with anterior cortical control compared to monolinguals. They showed attenuated responses for N2, theta and beta band rebound. Furthermore, they showed attenuated and later P3 responses as well as a smaller interference effect in P3 amplitudes.

Furthermore, our investigation into the effects of individual differences in bilingual experience onto neurocognitive and behavioural responses in the flanker task confirmed the attenuation of anterior control mechanisms with increasing level of bilingualism, as predicted by UBET framework, the BAPSS framework and the DRM. The increased duration of bilingual language usage was related to reduced bottom-up control, reflected in later P3 peaks and smaller increase in P3 amplitudes for interfering flankers. In addition, more intense and diverse bilingual language usage and increased language switching was related to reduced top-down control, reflected in a smaller midfrontal theta power effect. An effect of intensity and diversity of language usage on neural functioning is in line with assumptions of the ACH model (Abutalebi & Green, 2016; Green & Abutalebi, 2013), even if the ACH does not predict reduced anterior engagement.

The lack of predictive validity of our relative proficiency measure within the SEM analyses was not predicted (recall that it was dropped from SEM due to a high degree of covariance with intensity of use). However, this result stands in line with previous argumentation that, particularly past a certain point of proficiency, it is unlikely that higher proficiency in a language will *independently* contribute to neurocognitive adaptations beyond other experiential factors like degree of engagement with one’s languages. Recall that the relevant mechanisms for bilingualism-induced neurocognitive adaptation are active selection/inhibition of languages to facilitate communication. Thus, one’s competence in a language may only matter insofar as having the relevant representations to manage rather than how adherent to a target-like standard it is (for further discussion, see DeLuca et al., 2019). Thus, the more predictive measure of bilingual experience to affect different neurocognitive adaptations would be the extent to which the languages are engaged with, over time. Furthermore, greater degrees of engagement with one’s languages are likely to correlate with proficiency in them. Further research is needed, however, to assess this interpretation.

Maybe most interesting, modulations of bottom-up (i.e., stimulus-driven ERPs) and top-down mechanisms (induced activity, especially theta, see, e.g., Cavanagh & Frank, 2014; Helfrich & Knight, 2016) in the flanker task had opposite effects on the behavioural interference effect. Smaller interference effects were related to increased bottom-up control (i.e., larger P3 amplitudes and shorter latencies) on the one hand and reduced anterior top-down control (i.e., reduced theta power; Almabruk, Iyer, Tan, Roberts & Anderson, 2015; Jannsens, De Loof, Boehler, Pourtois, & Verguts, 2018; Mazaheri et al., 2014; Suzuki et al, 2018; Zavala et al, 2016) on the other hand. Together with the relationships of the bilingual experience factors with bottom-up and top-down control, these findings mean that longer bilingual duration effectively led to a larger interference effect, while increased intensity/diversity of bilingual usage and increased language switching led to a smaller interference effect. This is important in order to interpret the absence of a behavioural difference in interference suppression in our group comparison. It seems that the advantage of bilingual attenuation of frontal top-down control and the disadvantage of bilingual bottom-up control cancelled each other out.

The results of the flanker task and Simon task showed qualitive differences in control mechanisms. The Simon task, in contrast to the flanker task, showed no condition differences in alpha and beta band power. Furthermore, neuro-functional group differences present for the flanker task were largely absent. The only common result was attenuated anterior N2 responses in bilinguals, suggesting again a reduced anterior activity. In contrast to the flanker task, though, there was a) no group difference with regards to the interference effect in P3 amplitude, and b) bilinguals showed a larger instead of smaller interference effect in theta than monolinguals. While the latter could be interpreted as particularly strong bilingual engagement of anterior control during interference suppression, the widespread location of the effect suggests that control mechanisms were not a mere reflection of anterior cortical control. This stands in contrast to the particularly anterior location of the theta effect in the flanker task. Furthermore, in the Simon task, individual variations in bilingual experience were not related to neurocognitive reflections of bottom-up or top-down control, and the latter were not related to behavioural interference suppression. This stands in stark contrast to the findings for the flanker task.

We propose two possible reasons for the differences between tasks. Firstly, it may be the case that differences in task parameters led to differing degrees of control demands. In particular, the differing ratio of congruent-incongruent trials (50-50 in the Simon versus 75-25 in the flanker task) and the impact this has on monitoring demands (e.g., Costa et al., 2009) may have contributed to the discrepant findings. This is consistent with previous findings that bilingualism-induced behavioural performance effects may only be observed in tasks requiring a high degree of attentional control (Bialystok & Craik, 2022; Bialystok et al., 2004; Linck et al., 2008; Morales, Calvo, & Bialystok, 2013). Thus, while the paradigms used here were chosen for the purpose of our study because they had revealed superior performance for bilingual compared to monolingual participants, future studies might want to implement paradigms that are more demanding on attentional or cognitive control.

However, differences in interference effects on neural measures and the reversal of the group difference in interference-induced theta power between the tasks suggest that the flanker task was not simply more demanding. Instead, these results are in line with previous conclusions (e.g., Pratte, Rouder, Morey & Fend, 2013; Mansfield, van der Molen, Falkenstein & van Boxtel, 2013, but see Burle, Spieser, Servant & Hasbroucq, 2014) that the tasks recruit functionally distinct control mechanisms. These might be differentially impacted by bilingual experience. One indication for functional control differences is that congruency effect in flanker tasks increases with response time, whereas it decreases in the Simon task (Hübner & Töbel, 2019). Furthermore, asymmetries in the Simon task are particularly affected by lateralized spatial biases and not as much by response selection biases (Spironelli, Tagliabue & Umiltà, 2009), suggesting engagement of the orienting network is a particular important contributor to the interference effect. However, orienting network efficiency does not appear to differ between mono- and bilinguals (Costa et al, 2008; Costa et al, 2009; Hernández, Costa, Fuentes, Vivas & Sebastián-Gallés, 2010). It is thus perhaps not surprising that the Simon task is less sensitive to individual differences in bilingual experience than the flanker task. Future research in this area ought to utilise tasks which can assess multiple attentional networks (e.g., Attention network task, Fan et al., 2002) to more precisely delineate the effects of individual differences in bilingual experience on various aspects of attentional control.

Differences between bilingual and monolingual performance on the Simon task in previous studies has been inconsistent (e.g., Bialystok et al., 2004; among others). As suggested by Paap et al. (2015), it needs to be established under which circumstances group differences occur. One factor may be how bilingualism is operationalised. A recent study by Champoux-Larsson and Dylman (2021) demonstrates how this might change patterns of task performance on task. They found that measures of non-native language use in social situations and the composite bilingualism score of the LSBQ, which takes into account all kinds of measures of bilingual experience, predicted behavioural performance on the Simon task (with higher scores on these factors associated with slower reaction times, conferring with the results of the current study), while non-native language use at home and code switching did not predict performance. Importantly, age of second language acquisition did not predict performance when used dichotomously (i.e., late vs. early), but did when used as a continuous variable. This finding, along with those of the current study, suggest that utilising measures which capture a wide range of bilingual experiences can enable observation of effects which may be obscured by the dichotomous approach typically used in previous research. Another contributory factor to the differences in task may be the demands of the specific task, or the specific underlying cognitive processes that the task taps in to (Bialystok and Craik, 2022), as discussed above.

Despite the differences between the two tasks, we also found that bilinguals responded more slowly in both tasks, without a difference in interference effect. While this result appears at odds with many previous findings in the literature (e.g., Bialystok et al., 2004; Costa et al., 2009; Luk et al., 2011; Tao et al., 2011), it has been reported before (Vivas et al., 2017; Markiewicz, Mazaheri, & Krott, 2022). Markiewicz et al. (2022) found that the slow responses were caused by long response distribution tails and that the length of the tails was negatively correlated with P3 amplitude. Similarly, we found P3 amplitudes to be attenuated for bilinguals in the present study. The slower responses in bilinguals thus appear to be caused by reduced bottom-up control in our bilingual sample as a whole. Based on these results alone, one might conclude that the bilingual participants in our sample had a disadvantage in cognitive control. While this might be true for the bilingual and monolingual samples as a whole, our individual differences analysis shows that the picture is much more complicated. As such, taking individual differences into account leads to more interesting insights into the effects of bilingualism than group comparisons, with the latter having led to seemingly incompatible findings Importantly, we found that bottom-up and top-down processes had opposite effects on the behavioural interference effect. Our results therefore emphasise that slower responses in interference suppression tasks should not merely be interpreted as less efficient processing and therefore as a disadvantage. Much more interesting is that they may indicate varying adaptations to control demands, such as heightened conflict monitoring (e.g., Calabria, Hernández, Martin, & Costa, 2011; Zhou & Krott, 2018), or goal maintenance (e.g., Gullifer & Titone, 2021), particularly when performance is associated with reduced reliance on (neuronally expensive) cortical control, as appears to be the case for the bilinguals in our study.

The current study also supports previous work arguing that it is not sufficient to establish whether bilinguals and monolinguals differ in terms of behaviour in cognitive control tasks, it is important to investigate neuro-functional differences in order to conclude whether bilinguals perform differently or not. We have also seen that there is no simple one-to-one mapping of neuro-functional differences and performance differences, and different measures of bilingual experience (duration of bilingual language use versus intensity/diversity and language switching) can affect different neural measures and lead to opposite patterns of behavioural effects (larger or smaller interference effect). This also means that the term ‘bilingual advantage’ in attentional or cognitive control, often used in the literature, is inappropriate (see also Poarch & Krott, 2019).

In summary, the present findings are, to our knowledge, the first demonstration of a direct link between individual differences in bilingual experience, brain function and behavioural performance of an interference suppression task. They provide direct evidence of neurocognitive adaptations resulting from intensity, diversity and duration of non-native language use, as well as language switching. These factors were significantly associated with neurophysiological measures which predict behavioural performance and suggest an increasingly automatic mechanism for interference suppression (i.e., reduced reliance on cortical control). In order to elucidate the nature of the hypothesised bilingual adaptation in cognitive control, future studies in this area ought to fully consider the multi-faceted nature of both bilingual experience, and inhibitory control.

## Materials and methods

Data, analysis scripts and experiment files described in this section can be found at https://osf.io/ab8u3/.

### Participants

We recruited young adults via advertisements on the campus of the University of Birmingham, through social media, and word of mouth. Study participants were paid a small fee as compensation for their time. Two-hundred and thirty-nine participants (163 female, mean age: 22.9 years, SD: 3.7) completed session 1 of the experiment and therefore the flanker task. This included 71 native English-speaking participants (48 female, mean age: 21.0 years, SD: 2.6) who reported limited knowledge of a second language (i.e., complete beginner or elementary status). These were classified as monolinguals for our group comparison. The remaining 168 participants (115 female, mean age: 23.7 years, SD: 3.8) reported to be fluent in two, but not more than two languages (i.e., advanced or fluent). These participants were classified as bilinguals for our group comparison. Of the session 1 sample, 208 participants (146 female, mean age: 22.9 years, SD: 3.7) returned for session 2 and therefore for the Simon task. This included 60 participants classified as monolinguals (41 female, mean age: 20.9 years, SD: 2.8) and 148 participants classified as bilinguals (105 female, mean age: 23.7 years, SD: 3.8). Monolingual and bilingual groups were matched on a measure of general intelligence (Standard Progressive Matrices; Raven, Raven & Court, 2004). They were also matched on socioeconomic status (SES), which we measured by parental education level due to the fact that the large majority of our participants were students and thus did not differ much in terms of education or income (see Table 3).

All participants were pre-screened in accordance with the study’s eligibility criteria; they were required to be right-handed, have normal or corrected-to-normal vision and no history of head injuries resulting in concussion or any condition for which neurological damage was a feature, e.g., epilepsy. The study was conducted following the guidelines of the British Psychological Society code of ethics and approved by the Science, Technology, Engineering, and Mathematics (STEM) Ethical Review Committee for the University of Birmingham.

**Table 2:** Factor loadings for latent variables using CFA.

| Latent variables and their measures | Factor loadings |
| --- | --- |
| *Duration of bilingual experience* |  |
| Age of acquisition minus periods of non-use | 1.0 |
| *Intensity/diversity* |  |
| Language entropy | 0.868 |
| Non-English/English language use ratio (visual) | 0.820 |
| Non-English/English language use ratio (audio) | 0.734 |
| Please indicate which language(s) you generally use when speaking to: |  |
| - Friends | 0.816 |
| - Family (composite of use with parents, siblings, grandparents and other relatives) | 0.903 |
| Please indicate which language(s) you generally use for the following activities: |  |
| - Texting | 0.784 |
| - Emailing | 0.640 |
| - Watching TV | 0.603 |
| - Watching movies | 0.477 |
| - Browsing internet | 0.699 |
| - Reading | 0.72 |
| - Extracurricular activities | 0.557 |
| - Shopping/restaurants/other commercial services | 0.425 |
| Please indicate which language(s) you generally use for the following situations: |  |
| - Home |  |
| - Social activities | 0.613 |
|  | 0.756 |
| *Language switching* |  |
| How often do you switch languages between sentences at (home/studies/ work/other)? | 0.833 |
| How often do you switch languages within sentences at (home/studies/ work/other)? | 0.874 |
| How often does it happen that you switch between languages at (home/studies/work/other)? | 0.907 |
| I do not realize when I switch languages during a conversation at (home/studies/work/other)? | 0.539 |
| It is difficult for me to control the language switches I introduce during a conversation at (home/studies/work/other)? | 0.618 |
| When I switch languages in general at _____ I do it consciously (home/studies/work/other)? | 0.750 |
| I tend to consciously switch languages during a conversation at (home/studies/work/other) | 0.671 |
| There are situations in which I always choose to switch between the two languages at (home/studies/work/other) | 0.777 |
| There are certain topics or issues for which I normally choose to switch between the two languages at (home/studies/work/other) | 0.796 |
| Please indicate how often you engage in language switching |  |
| - With parents and family | 0.519 |
| - With friends | 0.704 |
| - On social media | 0.571 |
| *Relative proficiency* |  |
| Non-English/English self-reported proficiency ratio (visual) | 0.973 |
| Non-English/English self-reported proficiency ratio (audio) | 0.944 |
| Oxford (Quick) Placement Test | -0.571 |
| National Adult Reading Test | -0.412 |

**Table 3:** Mean SES and Raven’s Progressive Matrices scores. SES was assessed using the mean parental level of education where 0 = “No high-school diploma”, 0.25 = “High school diploma”, 0.5 = “Some post-secondary education”, 0.75 = “Post-secondary degree or diploma, 1 = “Graduate or professional degree”. Raven’s Progressive Matrices scores indicate the number of correct responses out of 60. Independent samples t-tests revealed no significant difference between the groups in either measure (all ps > 0.05)

| Group | SES (SD) | Raven's Progressive Matrices |
| --- | --- | --- |
| Monolinguals | 0.68 (0.28) | 44.5 (6.3) |
| Bilinguals | 0.66 (0.32) | 45.2 (7.2) |

### Procedure

The experiment comprised of the two cognitive control paradigms (detailed further in the Cognitive control paradigm section) and a comprehensive set of questionnaires (detailed further in the Bilingualism questionnaires section) to gather information about participants’ bilingual experience. These were administered over 2 sessions. Each session started with participants being measured for and fitted with an elasticated cap for EEG recordings, whilst they completed questionnaires. In session 1, participants filled in the Edinburgh Handedness Inventory (Oldfield, 1971), to confirm right-handedness, and the Language and Social Background Questionnaire (LSBQ). This was followed by the flanker task during which EEG was recorded. Subsequently, they completed the Oxford Quick Placement Test (OQPT) and a phonemic and a semantic verbal fluency (VF) task. In session 2, participants filled in the Switching Experience & Environments Questionnaire (SEEQ) and subsequently took part in the Simon task while EEG was recorded. They then completed the Standardised Progressive Matrices assessment (Raven, Raven & Court, 2004), second phonemic and semantic VF tasks, and finally the National Adult Reading Test (NART). In each session, participants completed one phonemic and one semantic VF task. Monolingual participants completed all VF tasks in English. Bilingual participants completed one phonemic and one semantic VF task each in their first and second languages, with languages counterbalanced between sessions.

Additionally, participants completed a resting state recording (7 minutes) as well as colour-shape and number-letter switching tasks. These were not part of the present research aims and relevant to a separate investigation.

#### Cognitive control paradigms

##### Simon (Simon & Wolf, 1963) and flanker (Eriksen & Eriksen, 1974) tasks

For the flanker and Simon tasks, we adopted the versions by Eriksen and Eriksen (1974) and Simon and Wolf (1963), respectively. The stimuli of the flanker task consisted of a row of five arrows presented either above or below a central fixation cross. Stimuli of the Simon task consisted of a red or blue square presented at either the left or right side of the display.

Figure 5 shows a schematic outline of the two tasks as well as the procedure for each testing session. For both tasks, each trial started with a 1500ms central fixation cross on a computer monitor. The stimulus was then presented and both stimulus and fixation cross remained on screen for 2000ms or until response. This was followed by a blank screen for 150ms. Interstimulus intervals (ISI) randomly varied between 1190 and 2179ms, with a mean ISI of 1654.6ms. Flanker stimuli occupied approximately 2° of visual angle and were presented approximately 0.72° (∼1cm) either above or below the fixation cross with equal probability. Simon stimuli occupied approximately 6° of visual angle and were presented approximately 5° (∼7cm) to the left or right of the fixation cross.

Each task began with a practice block of 24 trials (a mixture of 12 congruent and 12 incongruent trials). Participants indicated the direction of the central arrow (flanker task) or the colour of the stimuli (Simon task) using a Cedrus RB-834 response pad (Cedrus, USA). In the flanker task, they pressed a left button for left-pointing target arrows and a right button for right-pointing target arrows. In the Simon task, they pressed a left button for red squares and a right button for blue squares. The Flanker task consisted of 5 blocks of 96 trials each (72 congruent, 24 incongruent; total 480 trials), while the Simon task consisted of 7 blocks of 64 trials each (32 congruent, 32 incongruent; total 448 trials). The 25-75 ratio in the Flanker task was chosen because it is more taxing in terms of monitoring demands and therefore attentional control and has been found to distinguish more strongly between monolingual and bilingual speakers (see e.g., Costa et al., 2009; for other studies using this ratio, see, e.g., Zhou & Krott, 2018; Markiewicz, Mazaheri, & Krott, 2023).

Twelve participants’ behavioural data for the flanker, and 1 participant’s behavioural data for the Simon task, were lost due to corrupted or missing data files. Due to low commission of errors (2.2% for the Flanker and 3.5% for the Simon task), we did not analyse error rates. For reaction times (RTs), we analysed correct responses, excluding RTs < 200ms and > 2.5SD from participant means using the *trimr* package for R. A total of 4.6% of flanker trials and 6.2% of Simon trials were discarded from the analyses^3^.

**Figure 5:**
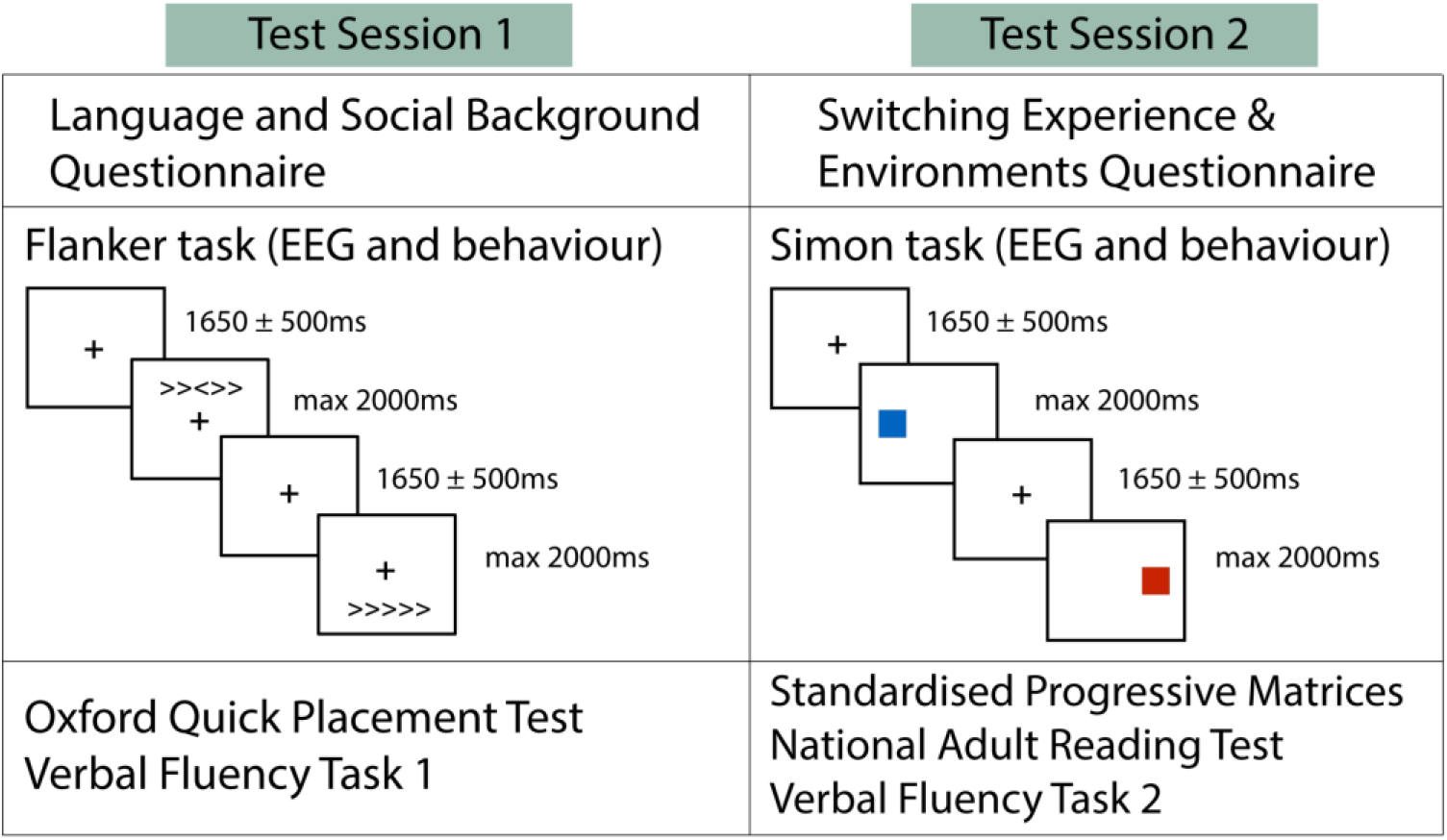
Testing procedure for study sessions as well as task parameters of the two EEG experiments.

#### Bilingualism questionnaires

##### Language and Social Background Questionnaire (LSBQ) (Anderson, Mak, Keyvani Chahi & Bialystok, 2018)

This questionnaire gathers information about patterns of language use of adults living in diverse communities where English is the dominant language. The questionnaire focuses on the extent of non-English proficiency and language use, both at home and socially.

Information from this questionnaire was entered into our SEM analysis (see Confirmatory Factor Analysis and Table 2 below).

##### Switching Experience and Environments Questionnaire (SEEQ; adopted from Hartanto & Yang, 2016, and Rodriguez-Fornells, Kramer, Lorenzo-Seva, Festman & Münte al., 2012)

This questionnaire assessed amount and type of language switching behaviour. It combined the Bilingual Switching Questionnaire (BSWQ; Rodriguez-Fornells et al., 2012) and the Code-Switching and Interactional Contexts Questionnaire (CSICQ; Hartanto & Yang, 2016). It tested 9 aspects of language switching and asked about typical time spent in 4 different environments (home, study, work and other, see section ‘switching’ in Table 2). Questions 1-4 are taken from the BSWQ, questions 8 and 9 from the CSICQ. Questions 5-7 are based on Green and Abutalebi’s (2013) distinction of interactional contexts of language use. Participants answered the questions for 4 different environments (home, study, work and other). Responses to each of the nine questions were adjusted for the percentage of time spent in each environment and were fed into our SEM analysis.

##### Oxford Quick Placement Test (OQPT; Syndicate, U. C. L. E., 2001)

The OQPT is a short version of the Oxford Placement Test and measures objective English proficiency. It contains 60 multiple choice items which assess the respondents’ knowledge of grammatical forms and vocabulary. The total number of correct answers were used for the SEM analysis.

##### Verbal Fluency (VF) tasks

As a second measure of objective language proficiency, we administered verbal fluency (VF) tasks. Participants were given 1 minute to list as many words as they could according to a certain phoneme (words beginning with ‘S’ or ‘F’ sound) or semantic (‘animals’ or ‘food’) criterion. Bilingual participants performed each task in L1/L2, while monolingual participants performed both tasks in L1.

##### National Adult Reading Test (NART; Nelson & Willison, 1991)

The third objective language measure was the NART, which assesses English vocabulary. It comprises 50 English words of increasing complexity, all of which are irregular in terms of rules of English pronunciation. Participants are asked to read the words. Correct pronunciation cannot be determined by phonetically decoding the word and thus relies on the knowledge of the words. Number of correctly pronounced words were used for the SEM analysis.

##### Standardised Progressive Matrices (Raven, Raven & Court, 2004)

This test is a general IQ measure and assesses nonverbal abstract reasoning. We conducted the test in a timed fashion, with participants having up to 25 minutes for completion. The number of correct answers were taken as the final score. Scores were used to compare monolingual and bilingual groups on general IQ.

### EEG Acquisition

Continuous electrophysiological recordings were obtained using a 64-channel eegoSports system (ANT Neuro, Enschede, The Netherlands) with Ag/AgCl passive electrodes positioned in a 10-10 system and implemented in elastic sensor caps (Waveguard classic, Advanced Neuro Technology B.V., Enschede, Netherlands). The EEG was acquired with online reference to the CPz channel, at a sampling rate of 500 Hz, with a 30-Hz low-pass filter (24 dB/octave) and a 0.05-Hz high-pass filter, implemented in the EEGosports firmware. Impedances were kept below 20kOhms. Additional bipolar electrodes were placed beside each eye for horizontal and above/below the right eye for vertical electro-oculogram recordings.

### EEG Pre-processing & Analysis

Offline EEG recordings were imported into EEGLAB (Delorme & Makeig, 2004) for visual examination, and excessively noisy or flat channels were removed. EEG recordings were then high-pass filtered at 1Hz for the purpose of conducting independent components analysis (ICA), as it has been shown that this results in significantly improved signal-to-noise ratio and classification accuracy of components relative to not filtering (Winkler, Debener, Muller & Tangermann, 2015). ICA weights obtained from the filtered data were then added back to the original data. The resulting data were re-referenced to an average montage, and segmented around stimulus onset, with a 2000ms baseline and epochs extending to 3000ms post-stimulus. Blinks, eye movements and muscle artifact components were then identified and removed, with an average of 2.22 components removed from the data for the flanker task (ML: 2.04, BL: 2.27), and 2.75 components removed for the Simon task. (ML: 2.72, BL: 2.76). We rejected trials if responses were incorrect, RTs < 200ms or >2.5SD of the mean for each condition per participant. This led to 2.12% of monolinguals’ trials (6.96% incongruent, 0.5% congruent) and 1.46% of bilinguals’ trials (4.27% incongruent, 0.52% congruent) being discarded for the flanker task and 3.07% of monolinguals’ trials (3.77% incongruent, 2.36% congruent) and 2.79% of bilinguals’ trials (3.38% incongruent, 2.19% congruent) for the Simon task. Visual inspection of the data was then conducted to identify and remove epochs which still contained excessive noise or other artifacts. In total, 9.83% of flanker trials were rejected, leaving an average of 432.8 trials per participant (ML: 429, BL: 434.5). 10.19% of Simon trials were rejected, leaving an average of 402.4 trials per participant (ML: 402.9, BL: 402.1). Subsequently, channels that were originally removed due to excessive noise were interpolated.

Time-frequency representations (TFRs) of power were estimated on the EEG epochs with the FieldTrip “mtmconvol” method and using an adaptive sliding time window (Hanning taper) with the length of 3 cycles per each frequency of interest. This approach has been taken previously by Mazaheri et al. (2009), Poulisse et al. (2020), and van Diepen et al. (2015). The analysis included the frequency of interest of 2–30 Hz in steps of 1 Hz, and the time of interest of −1 to 2 s in steps of 0.05 s.

Time-locked averaged ERPs were subtracted from the power spectra for each individual and condition in order to identify induced oscillatory activity in isolation from evoked activity (Cohen, 2014; Sauseng et al, 2009, Mazaheri and Picton, 2005). To examine neural correlates of inhibitory control, power spectra obtained from congruent trials were subtracted from those obtained from incongruent trials (Cohen & Ridderinkhof, 2013; Nigbur, Ivanova & Stürmer, 2011). These difference spectra formed the basis of the frequency analyses and structural equation modelling (SEM) reported below.

### Confirmatory Factor Analysis

Responses of questionnaires (LSBQ, SEEQ) and proficiency measures (O-QPT, NART & VF tasks) were loaded onto 4 latent variables to test the effect of bilingual experience on interference suppression (derived from the UBET model; DeLuca et al, 2020). These were *duration of bilingual language use*, *intensity and diversity of language use* (hereafter *intensity/diversity)*, *language switching* and *relative proficiency*.

The latent variables were composed by confirmatory factor analysis (CFA) using the package *lavaan* (Rosseel, 2012) in R (version 4.1.2; R Core Team, 2021). Table 2 lists the measures that were loaded onto the different variables together with their factor loadings.

*Duration of bilingual language use* was calculated by subtracting periods of second language non-use from an overall measure of second language duration (age at time of testing minus age of acquisition).

The factor *intensity/diversity* captured patterns of language use across different contexts and was based on responses from the LSBQ (Anderson et al, 2018) and a measure of language entropy. More specifically, it assessed the ratio of first language/second language usage across different interaction partners (friends, family), activities (e.g., texting, emailing, browsing internet etc.) and contexts (home, social activities). Language entropy was calculated using the *language entropy* function implemented in R (Gullifer & Titone, 2018).

The *language switching* factor was based on the SEEQ and assessed language switches in terms of frequency and type (e.g., conscious, unconscious) and across different contexts (e.g., with friends, family and on social media).

The factor *relative proficiency* was based on scores from the Oxford Placement Test (Quick version; O-QPT) and National Adult Reading Test (NART), along with a ratio of second language-English verbal fluency, as measured by the verbal fluency tasks.

In general, error variances were correlated where items were theoretically proposed to overlap (e.g., language used while watching movies and TV, or language used while emailing and browsing the internet etc.)

### Structural Equation Modelling

We hypothesised that the latent variables derived from the CFA described above (i.e., *intensity/diversity, language switching, duration of bilingual language use* and *relative proficiency*) would be associated with functional neurocognitive adaptations, which in turn would predict the behavioural effects of interference during Flanker and Simon task performance. We conducted structural equation modelling (SEM) using the *lavaan* package in R (Rosseel et al, 2012) to examine these hypotheses.

We included only participants with full datasets in this analysis. Reasons for missing data was primarily questionnaire data that was not submitted or stored correctly, corrupt behavioural data files or unusable EEG data from either task (due to low recording quality, missing stimuli tags etc.) Therefore, the analysis reported in this section is conducted on a dataset of 186 participants (135 categorised as bilinguals).

The latent language variables obtained from CFA were loaded onto two latent factors which corresponded to theoretically derived variables of interest (DeLuca et al., 2020): *Intensity/diversity* and *language switching* were loaded onto a factor labelled *executive function demands (EF demands)*, while *relative proficiency* and *duration of bilingual language use* were loaded onto a factor labelled *efficiency.* A covariance term was included for *EF demands* and *efficiency.* CFA revealed that observed variables loaded significantly onto these latent variables, but negative variances were observed for *relative proficiency* and *intensity/diversity*. Examination of the correlation matrix revealed this was likely caused by collinearity between these two variables (r > 0.95). *Intensity diversity* was considered the more robust measure of the two as it was loaded by a larger number of observed variables and its estimated variance was less negative. Thus, *relative proficiency* was removed from the model and *duration of bilingual language use* was set as a single-indicator variable.

Functional neurocognitive adaptations were divided into bottom-up control and top-down control. Interference effects for P300 and N2 measures were loaded onto the latent variable *bottom-up control*, while interference effects for theta and alpha power were loaded onto the latent variable *top-down control*. Spectral power was extracted from regions-of-interest (ROIs) and timepoints-of-interest (TOIs) which corresponded to the maximal differences between conditions, as identified by the cluster-based permutation tests reported above. Thus, induced theta power was extracted from 250-750ms post-stimulus for the flanker task, and 100-500ms post-stimulus for the Simon task, from a ROI comprised of several frontocentral electrode sites (FCz, Fz, FC1, FC2, F1 & F2). Induced alpha power was extracted from 600-1000ms post-stimulus from a ROI of occipital regions (O1, O2 & Oz). ERP amplitude and latency measures corresponded to the condition difference (i.e., incongruent minus congruent) and were extracted from the same electrode sites as used in the ANOVA described above.

The RT measure represented the RT difference between incongruent and congruent correct trials, as determined by the procedure described above.

## Acknowledgements and funding sources

This study was funded through a research grant (ES/R005311/1) from the Economic and Social Research Council (United Kingdom). We thank Consuelo Vidal Gran, Samantha Millard, and Zlatomira Ilchosvka for their help with participant recruitment and testing as well as the participants for taking part in this multi-session experiment.

## Footnotes

1 Including mean parental education level (as a proxy measure of SES) and Raven’s Progressive Matrices scores (as a proxy measure of IQ) as covariates in our linear mixed effects model of the behavioural data did not change the pattern of results. There was still an effect of language group (ML/BL) on RTs in the Flanker task: t(198.1) = 2.457, p = 0.0149, and a trend for such an effect on the Simon task: t(198.28) = 1.786, p = 0.076. Importantly, including the covariates did not change the non-significant interaction of group and condition (flanker task: t(196.5) = -0.658, p = 0.5114; Simon task: t(196.3) = -0.007, p = 0.994). Furthermore, neither of the two covariates was significantly associated with behavioral performance in either task (p > 0.05, and groups did not significantly differ on these factors.

2 Visual inspection suggested a group difference in P3 amplitude. However, a cluster-based permutation test in a 0-1000ms window showed no significant difference.

3 We repeated the behavioral analyses using untrimmed data and this did not alter the pattern of findings.

